# SARS-CoV-2 Omicron neutralization by therapeutic antibodies, convalescent sera, and post-mRNA vaccine booster

**DOI:** 10.1101/2021.12.22.473880

**Authors:** Sabrina Lusvarghi, Simon D. Pollett, Sabari Nath Neerukonda, Wei Wang, Richard Wang, Russell Vassell, Nusrat J. Epsi, Anthony C Fries, Brian K Agan, David A. Lindholm, Christopher J. Colombo, Rupal Mody, Evan C. Ewers, Tahaniyat Lalani, Anuradha Ganesan, Emilie Goguet, Monique Hollis-Perry, Si’Ana A. Coggins, Mark P. Simons, Leah C. Katzelnick, Gregory Wang, David R. Tribble, Lisa Bentley, Ann E. Eakin, Christopher C. Broder, Karl J. Erlandson, Eric D. Laing, Timothy H. Burgess, Edward Mitre, Carol D. Weiss

## Abstract

The rapid spread of the highly contagious Omicron variant of SARS-CoV-2 along with its high number of mutations in the spike gene has raised alarm about the effectiveness of current medical countermeasures. To address this concern, we measured neutralizing antibodies against Omicron in three important settings: (1) post-vaccination sera after two and three immunizations with the Pfizer/BNT162b2 vaccine, (2) convalescent sera from unvaccinated individuals infected by different variants, and (3) clinical-stage therapeutic antibodies. Using a pseudovirus neutralization assay, we found that titers against Omicron were low or undetectable after two immunizations and in most convalescent sera. A booster vaccination significantly increased titers against Omicron to levels comparable to those seen against the ancestral (D614G) variant after two immunizations. Neither age nor sex were associated with differences in post-vaccination antibody responses. Only three of 24 therapeutic antibodies tested retained their full potency against Omicron and high-level resistance was seen against fifteen. These findings underscore the potential benefit of booster mRNA vaccines for protection against Omicron and the need for additional therapeutic antibodies that are more robust to highly mutated variants.

**One Sentence Summary:** Third dose of Pfizer/BioNTech COVID-19 vaccine significantly boosts neutralizing antibodies to the Omicron variant compared to a second dose, while neutralization of Omicron by convalescent sera, two-dose vaccine-elicited sera, or therapeutic antibodies is variable and often low.

## INTRODUCTION

In November 2021 a new SARS-CoV-2 variant, named Omicron (Pango lineage B.1.1.529), was identified as a variant of concern (VOC). Its rapid spread in Africa and unusually high number of mutations, especially in the spike gene, has triggered intense international efforts to track the variant’s spread and evaluate its effects on the potency of therapeutics and vaccines. The predominant strain of Omicron has mutations in the spike gene encoding 15 amino acid changes in the receptor binding domain (RBD) of the spike surface protein (G339D, S371L, S373P, S375F, K417N, N440K, G446S, S477N, T478K, E484A, Q493R, G496S, Q498R, N501Y, and Y505H). The RBD mediates virus attachment to the ACE2 receptor on target cells and is the principal target of neutralizing antibodies that contribute to protection against SARS-CoV-2. Many of these RBD changes have been previously reported to reduce the effectiveness of several therapeutic neutralizing antibodies (reviewed in Corti et al(*1*)). A recent study reports that the full complement of RBD substitutions in the Omicron spike compromises the potency of over 85% of 247 anti-RBD monoclonal antibodies (mAbs) tested(*2*). Preliminary reports indicate substantial immune evasion to two-dose vaccine-elicited sera(*3-7*), booster-elicited sera(*8-16*), genotype-varying convalescent sera(*3, 5, 6*), and several mAbs(*2, 6*). However, study populations and methods vary widely among the studies to date, and many lack critical details about host characteristics. Moreover, studies have not examined how host demography predicts these neutralizing humoral responses, and examination of how infection by a broader diversity of SARS-CoV-2 Delta and non-Delta genotypes is important for further insights into how genetic diversity may correlate with cross-neutralizing antibody responses.

Here we used a pseudovirus neutralization assay(*16*) to measure antibody neutralization of SARS-CoV-2 Omicron in three important contexts: (1) antibodies induced after two or three doses of the Pfizer-BioNTech Covid-19 (Pfizer/BNT162b2 mRNA) vaccine, (2) antibodies induced from infection by different SARS-CoV-2 variants and (3) therapeutic antibodies under emergency use authorization (EUA) or in later stages of clinical development. We compared the magnitude of neutralization escape by Omicron to D614G (referred to as wild type, WT) and Delta SARS-CoV-2 variants to help inform public health decisions and offer further data toward correlate of protection research.

## RESULTS

### Three immunizations of the Pfizer/BNT162b2 mRNA COVID-19 vaccine significantly boosts neutralizing antibodies to the Omicron variant compared to two-vaccinations

The emergence of the Omicron variant coincided with recommendations for booster immunizations, particularly for at risk populations. We studied the neutralization titers of 39 generally healthy, adult healthcare workers participating in the Prospective Assessment of SARS-CoV-2 Seroconversion study (PASS study, Table 1)(*17*) who received the full primary series (1^st^ and 2^nd^) and booster (3^rd^) immunizations with the Pfizer/BNT162b2 vaccine. We chose to study sera at peak responses after the full primary series vaccination rather than after 6 months because 6-months titers are often very low(*10, 18*).

**Table 1.** Demographic data for participants receiving Pfizer/BNT162b2 initial vaccine series and booster.

|  | N (%) |
| --- | --- |
| **Sex** |  |
| Female | 25 (64.1) |
| Male | 14 (35.9) |
| **Race** |  |
| White | 26 (66.6%) |
| Asian | 8 (20.5%) |
| Black | 4 (10.3%) |
| Multiracial | 1 (2.6%) |
| **Occupation** |  |
| Nurse | 11 (28.2%) |
| Physician | 11 (28.2%) |
| Physical/Occupational/Recreational Therapist | 9 (23.1%) |
| Medical Technician | 3 (7.7%) |
| Lab Personnel | 3 (7.7%) |
| Social Worker | 1 (2.6%) |
| Psychologist | 1 (2.6%) |
| **Anti-N seroconversion after vaccination and before**<br>**boost** |  |
| Positive | 17 (43.6%) |
| Negative | 22 (56.3%) |
| **Age** |  |
| Mean age ± SD (range) | 45 ± 11 (26 - 69) |
| **Time between second vaccine and sample collection** |  |
| Mean days ± SD (range) | 30 ± 11 (28 - 34) |
| **Time between second vaccine and booster dose** |  |
| Mean days ± SD (range) | 267 ± 14 (218-310) |
| **Time between booster dose and sample collection** |  |
| Mean days ± SD (range) | 43 ± 17 (7 - 93) |

Neutralization activity against Omicron above the threshold of our assay (1:40 dilution) was only observed in 12.8% (5/39) of serum samples obtained at a mean of 30 ± 11 days after the 2^nd^ Pfizer/BNT162b2 vaccination (Figure 1A). After the 2^nd^ vaccination neutralization titers against Omicron (geometric mean titer, GMT 22) were 25.5-fold lower than neutralization titers against WT (GMT 562). By contrast, neuralization titers against Delta (GMT 292) were 1.9-fold lower than WT. Neutralization titers from the same individuals collected 43 ± 17 days after the 3^rd^ Pfizer/BNT162b2 vaccination were 8.9-fold greater against WT (GMT 5029) compared to titers after the 2^nd^ vaccination. The titers against Omicron after the 3^rd^ vaccination (GMT 700) showed 31.8-fold increase compared to titers after the 2^nd^ vaccination, whereas titers against Delta after the 3^rd^ vaccination (GMT 1673) were only 5.7-fold higher than the titers after the 2^nd^ vaccination. Importantly, all individuals had measurable neutralizing titers against Omicron after the 3^rd^ vaccination, highlighting the potential for increased protection by a booster vaccine.

**Fig. 1.**
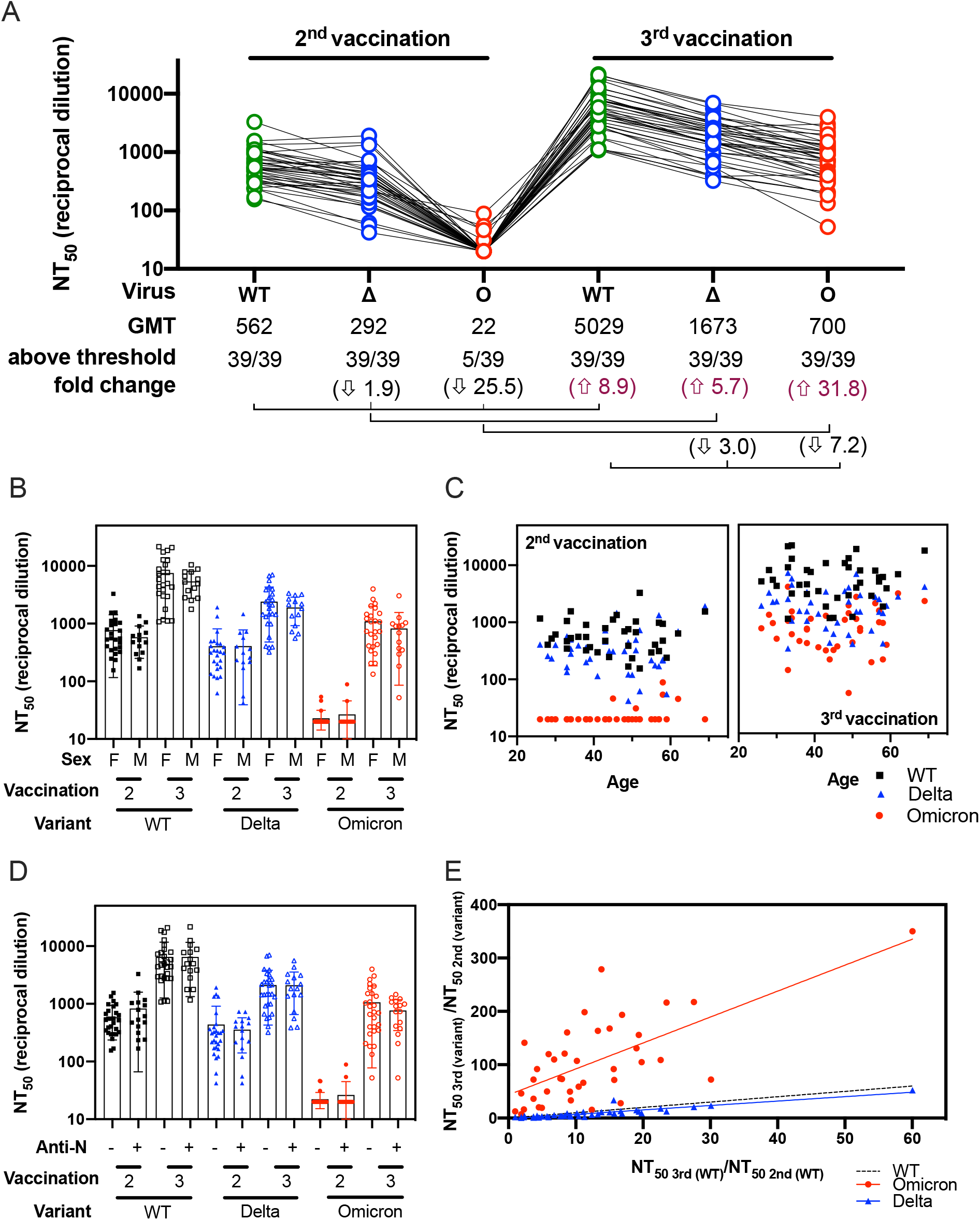
Sensitivity of the Omicron variant to neutralization by Pfizer/BNT162b2 vaccinee sera. (A) Neutralization assays were performed using lentiviral pseudoviruses bearing SARS-CoV-2 WT (B.1 lineage, D614G), Delta (Δ, B.1.617.2 lineage, T19R, G142D, E156-, F157-, R158G, L452R, T478K, D614G, P681R and D950N) and Omicron (O, B.1.1.529 lineage, A67V, del69-70, T95I, del142-144, Y145D, del211, L212I, ins214EPE, G339D, S371L, S373P, S375F, K417N, N440K, G446S, S477N, T478K, E484A, Q493R, G496S, Q498R, N501Y, Y505H, T547K, D614G, H655Y, N679K, P681H, N764K, D796Y, N856K, Q954H, N969K and L981F) with sera from 39 healthcare workers after immunization with two and three doses of Pfizer/BNT162b2 vaccine. Sera was obtained at a mean of 30 ± 11 days after immunization with two doses and at a mean of 43 ± 17 days after booster vaccination. Each serum was run in duplicate in two independent experiments against each pseudovirus to determine the 50% neutralization titer (NT_50_). The geometric means titers (GMT), the number of NT_50_ above threshold (1:40) and the fold change are indicated. Titers below 1:40 were set at 20 to calculate GMTs. Arrows indicate increase or decrease relative to WT. Connecting lines indicate serum from the same individual. The demographic information for this sera cohort is provided in Table 1. (B) NT_50_ by sex after 2^nd^ or 3^rd^ vaccination. (C) NT_50_ by age after 2^nd^ and 3^rd^ vaccination. (D) 2^nd^ or 3^rd^ vaccination NT_50_ according to anti-N (nucleocapsid protein) seroconversion between 2^nd^ and 3^rd^ vaccination. (E) The ratios between the neutralization titers after the 3^rd^ and the 2^nd^ immunization for Omicron and Delta were plotted against the corresponding ratios for WT. For panels B-E, black squares correspond to WT, blue triangles correspond to Delta and red circles correspond to Omicron.

We also evaluated how sex or age of the individual might affect the titers in sera after vaccination. We did not observe a trend according to sex or age after the 2^nd^ or 3^rd^ vaccination (Figure 1B and 1C, respectively). A total of 17/39 individuals had a positive anti-N (SARS-CoV-2 nucleoprotein) seroconversion during regular scheduled blood sampling between the 2^nd^ and the 3^rd^ immunization(*17*). No symptoms of COVID-19 were reported by the subjects, possibly indicative of silent infection or exposure to SARS-CoV-2 or other coronaviruses. We did not find any trends when comparing differences in neutralizing antibody titers between individuals who had anti-N seroconversion and those who did not (Figure 1D). To assess the breadth of neutralization responses against Omicron induced by boosting, we compared the change in titers against Omicron or Delta relative to WT after the 2^nd^ and 3^rd^ vaccination. To account for variability in the antibody titers from one individual to another, we calculated the ratio between the neutralization titers after the 3^rd^ and the 2^nd^ vaccination for each variant and plotted this ratio for Omicron and Delta against WT (Figure 1E). We observed that the ratios of NT_50_ titer from the 3^rd^ to the 2^nd^ vaccination for Omicron relative to the corresponding ratios for WT were higher than the ratios for Delta relative to WT, suggesting that the 3^rd^ vaccination broadened responses to the distantly-related variant.

### Neutralization of Omicron by convalescent sera from individuals infected by different variants shows lower neutralization titers compared to the infecting variant

To investigate the potency of infection-induced neutralizing antibodies against Omicron, we compared neutralization titers against WT, Delta, and Omicron in convalescent sera obtained at a mean of 30.2 ± 9.3 days post-symptom onset from unvaccinated individuals infected with WT, Alpha, Beta, or Delta variants who were enrolled in the Epidemiology, Immunology, and Clinical Characteristics of Emerging Infectious Diseases with Pandemic Potential (EPICC) study(*19*). Genotypes of the infecting variants were sequenced for all cases (Table 2 and Supplementary Table). These convalescent sera were complemented by a Beta-convalescent serum from another protocol.

**Table 2:**
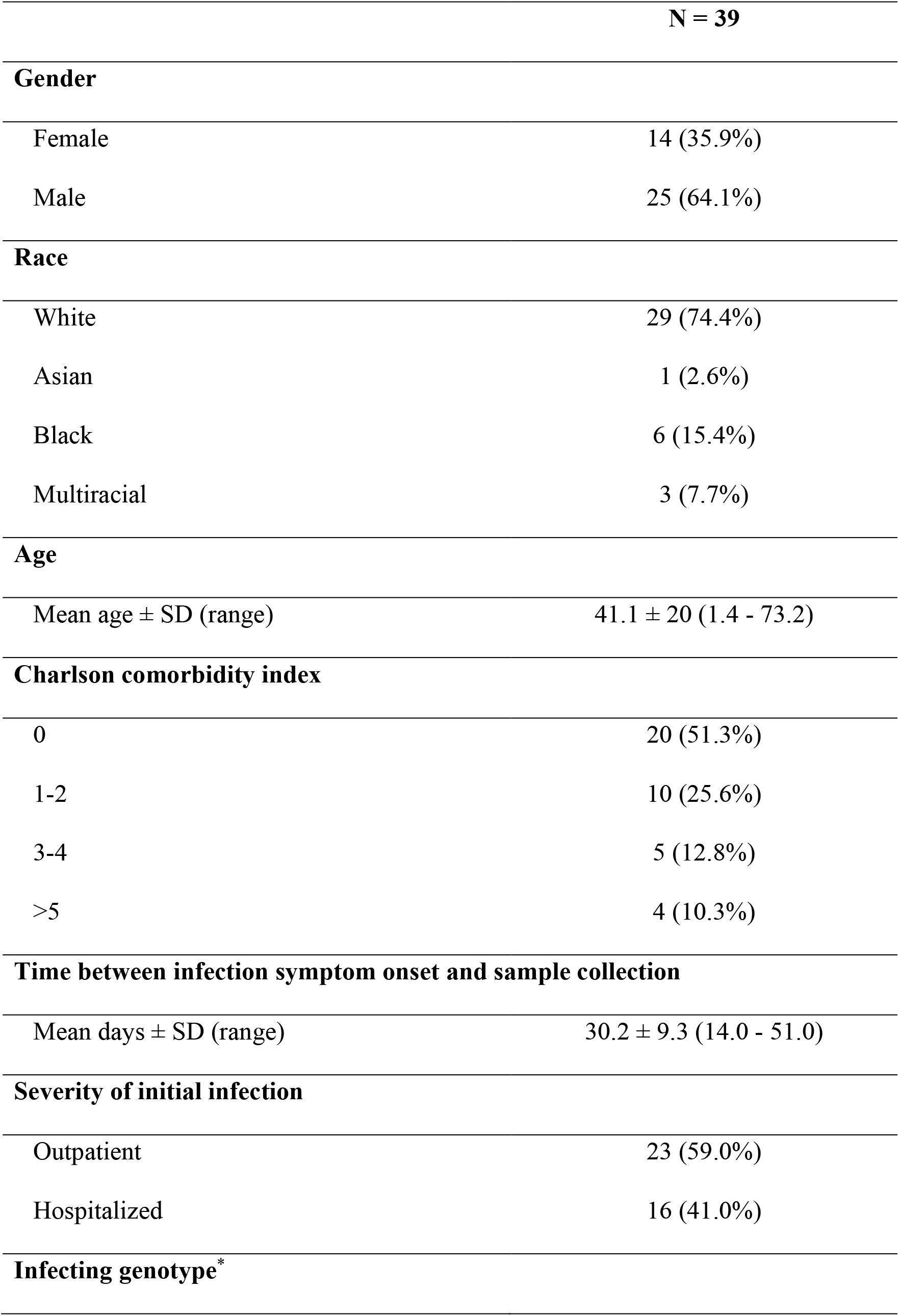
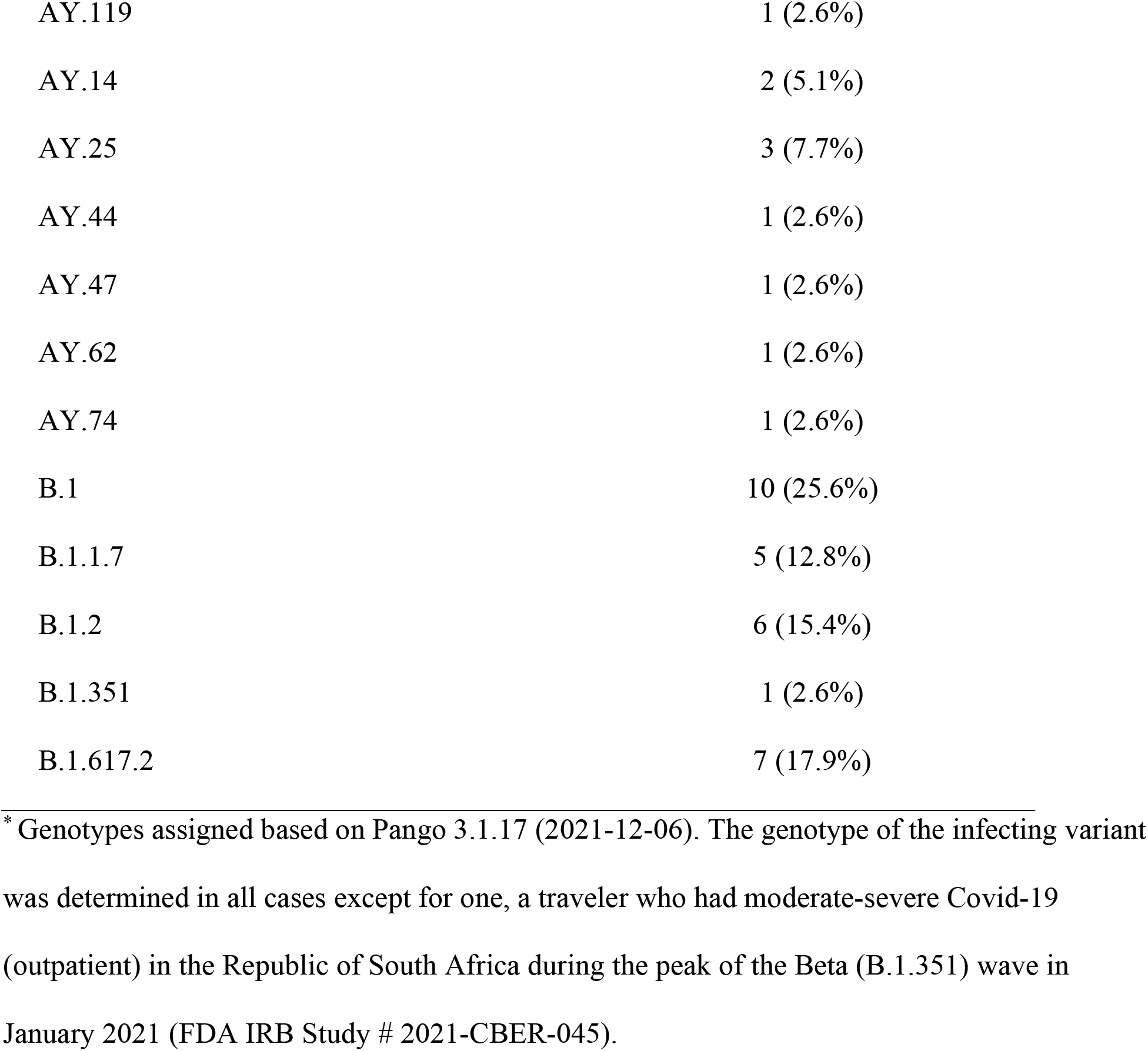
Characteristics of unvaccinated infections providing convalescent sera.

Figure 2 shows that the Delta variant was modestly more resistant to neutralization than WT by sera from most individuals infected with WT, Alpha, or Beta viruses (1.1 to 1.9-fold), while neutralization titers were much more reduced against Omicron (2.3 to 70.1-fold). A total of 2/10 and 0/20 individuals infected with WT (B.1 and B.1.2) variants, respectively, had a response above the threshold against Omicron, whereas 2/5 and 2/2 individuals infected with Alpha (B.1.1.7) and Beta (B.1.351) variants, respectively, were above threshold. Convalescent sera from individuals infected with Delta (B.1.617 and AY.14/.25/.44/.47/.62/.74/.119) variants generally had higher neutralization titers against Delta (2.8 to 13.5-fold) compared to WT, indicating more focused antibody responses to epitopes in the Delta spike. Convalescent sera from these Delta-infected individuals showed 22.1 to 74.4-fold lower neutralization titers against Omicron compared to Delta. However, a total of 6/7 individuals infected with the B.1.617.2 variant and 9/10 individuals infected with an AY variant had titers above threshold against Omicron.

**Fig. 2.**
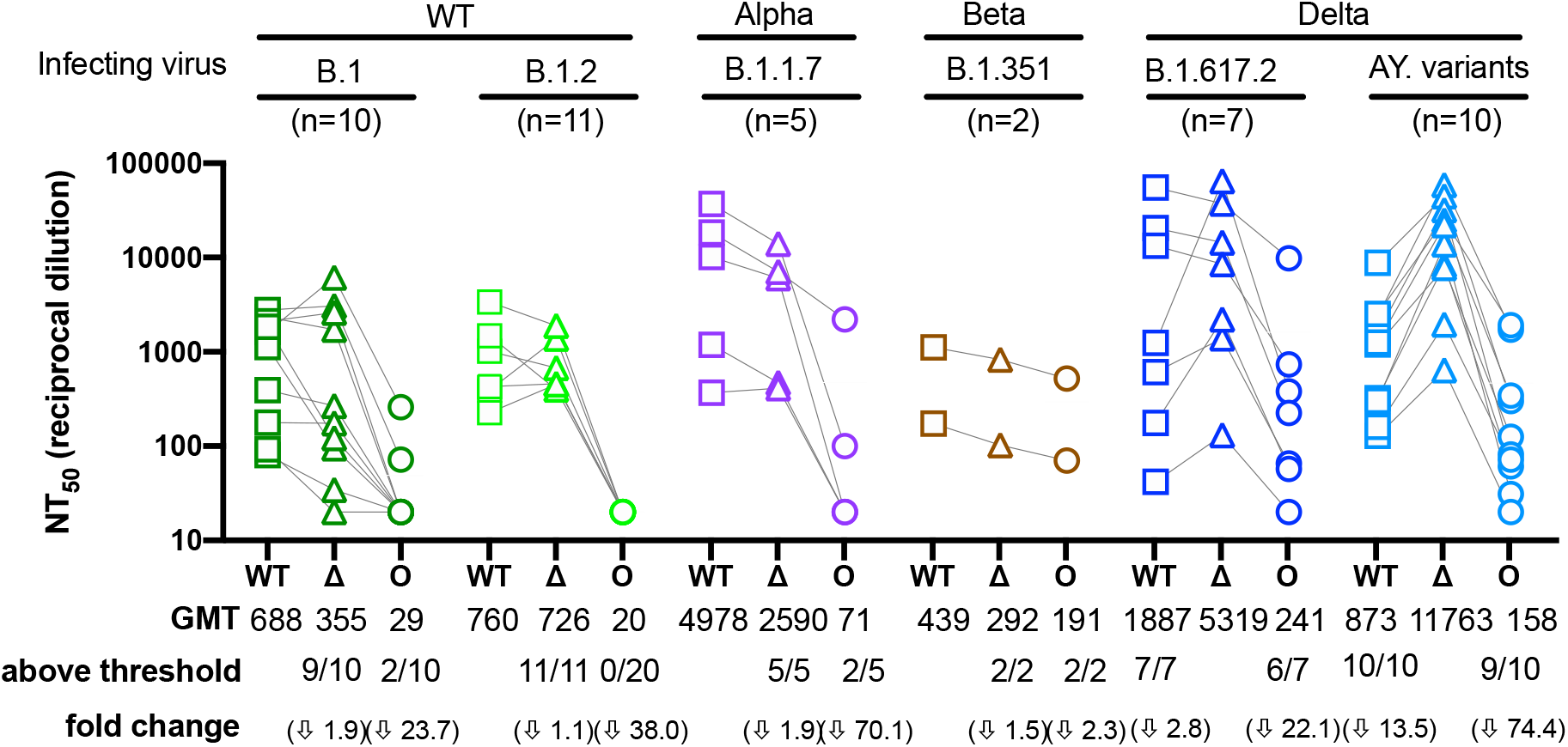
Sensitivity of the Omicron variant to neutralization by convalescent sera. Neutralization assays were performed using convalescent sera from persons infected with genotyped variants from B.1, B.1.2, B.1.1.7, B.1.351, B.1.617.2, AY.14, AY.25, AY.44, AY.47, AY.62, AY.74 or AY.119 lineages (Table 2 and Supplementary Table). Both B.1 and B.1.2 have no mutations in the receptor binding domain and were therefore considered WT, whereas some of the AY mutants have additional mutations in the RBD relative to B.1.617.2. Each serum was run in duplicate against WT, Delta, and Omicron to determine the NT_50_. The geometric means (GMT), the number of NT_50_ above threshold (1:40) and the fold change are indicated. Titers that did not inhibit at the lowest dilution tested (1:40) were assigned a titer of 20 for GMT calculations. Arrows indicate decrease relative to the infecting variant. Connecting lines indicate serum from the same individual. Data shown represent two independent experiments each with an intra-assay duplicate. Squares correspond to WT, triangles correspond to Delta, and circles correspond to Omicron.

### Boosting reduces apparent antigenic differences between WT and Omicron variants

We applied antigenic cartography to explore how the convalescent and post-vaccination sera distinguish the different spike antigens. Antigenic maps were made separately using neutralizing antibody titers from individuals infected with the different variants or from individuals after the 2^nd^ or 3^rd^ vaccination (Figure 3). Convalescent sera were more heterogeneous compared to the post-vaccination, with each convalescent serum clustering closer to the infecting variant, as expected. The 3^rd^ post-vaccination sera were more tightly clustered around WT compared to the 2^nd^ post-vaccination sera. The antigenic distances between Omicron and WT were large for all three sera sets, but the antigenic distance between Omicron and WT were smaller after the 3^rd^ vaccination (7.2-fold drop) compared to either convalescent or the 2^nd^ vaccination sera (49.4-fold drop and 39.4-fold drop, respectively), in agreement with the titer changes in Figure 1A. Small distance changes between WT and Delta were observed for all three sera sets (2.0 to 3.6-fold difference).

**Fig. 3.**
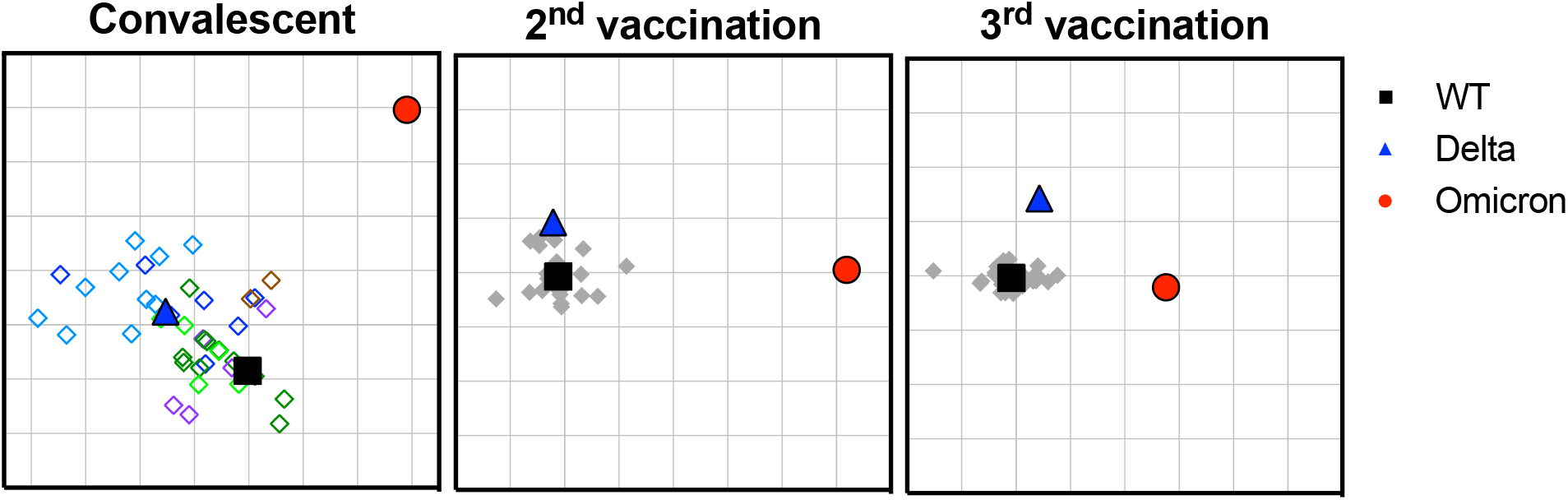
Antigenic cartography of convalescent and vaccinee sera against WT, Delta and Omicron. Antigenic maps were separately generated from convalescent (left panel), 2^nd^ vaccination (middle panel) or 3^rd^ vaccination (right panel) sera. Convalescent sera are shown in diamonds as follows: B.1 (dark green), B.1.2 (light green), B.1.1.7 (purple), B.1.351 (brown), AY variants (light blue), and B.1.617.2 (dark blue). Gray diamonds correspond to post-vaccination sera. Each grid square corresponds to 2-fold dilution in the neutralization assay. Black squares correspond to WT variant. Blue triangles correspond to Delta variant. Red circle corresponds to Omicron variant.

### The potency of most therapeutic antibody products is compromised against Omicron

As part of the US government COVID-19 response effort to speed development of therapeutics for COVID-19, we assessed the neutralization of Omicron by 24 therapeutic antibody products currently under EUA or in late stages of clinical development. This panel includes 15 single therapeutic neutralizing antibody products (nAbs), six combination nAbs (cnAbs) and three polyclonal antibody preparations (pAbs). The manufacturers provided these clinical products for side-by-side comparisons of potency against variants but required blind coding of these antibody products for publications. Previously, we reported that several single substitutions in the spike protein of other variants conferred resistance to some of these products(*20*), but a similar assessment has not been performed on the Omicron spike.

Figure 4A shows the neutralization curves for each product against WT and Omicron. To quantify the relative drop in nAb potency we calculated ratios between the 50% inhibitory concentration of Omicron to WT (Figure 4B). These ratios do not account for the absolute potencies of the nAbs but do allow for a uniform comparison of the changes in potency against Omicron relative to WT for all the antibody products. We note that the majority of successful clinical trials were performed when predominant strains had a ratio of near 1 compared to WT. However, absolute potency using IC_50_ ng/ml as a measure has not been established as a correlate of protection. We defined ratios between of 5 to 50 as a benchmark for partial or moderate resistance and ratios of > 50 as a benchmark for more complete resistance relative to WT. While the clinical relevance of the IC_50_ changes has not been determined for any *in vitro* assay, these cutoffs were chosen because the therapeutic levels of antibody therapeutics may be high enough to overcome low levels of resistance. By the fold-change measure, only three of 15 nAbs retained near full potency against Omicron compared to WT, and only one retained partial potency. Two cnAbs retained partial potency, while the remaining four cnAbs showed complete loss of neutralization potency. All three pAbs showed reduced neutralization potency (ratios 13-17) against Omicron, in agreement with the data from convalescent and vaccinated individuals. These findings raise concerns that many of the available therapeutic antibody products may not be effective against Omicron.

**Fig. 4.**
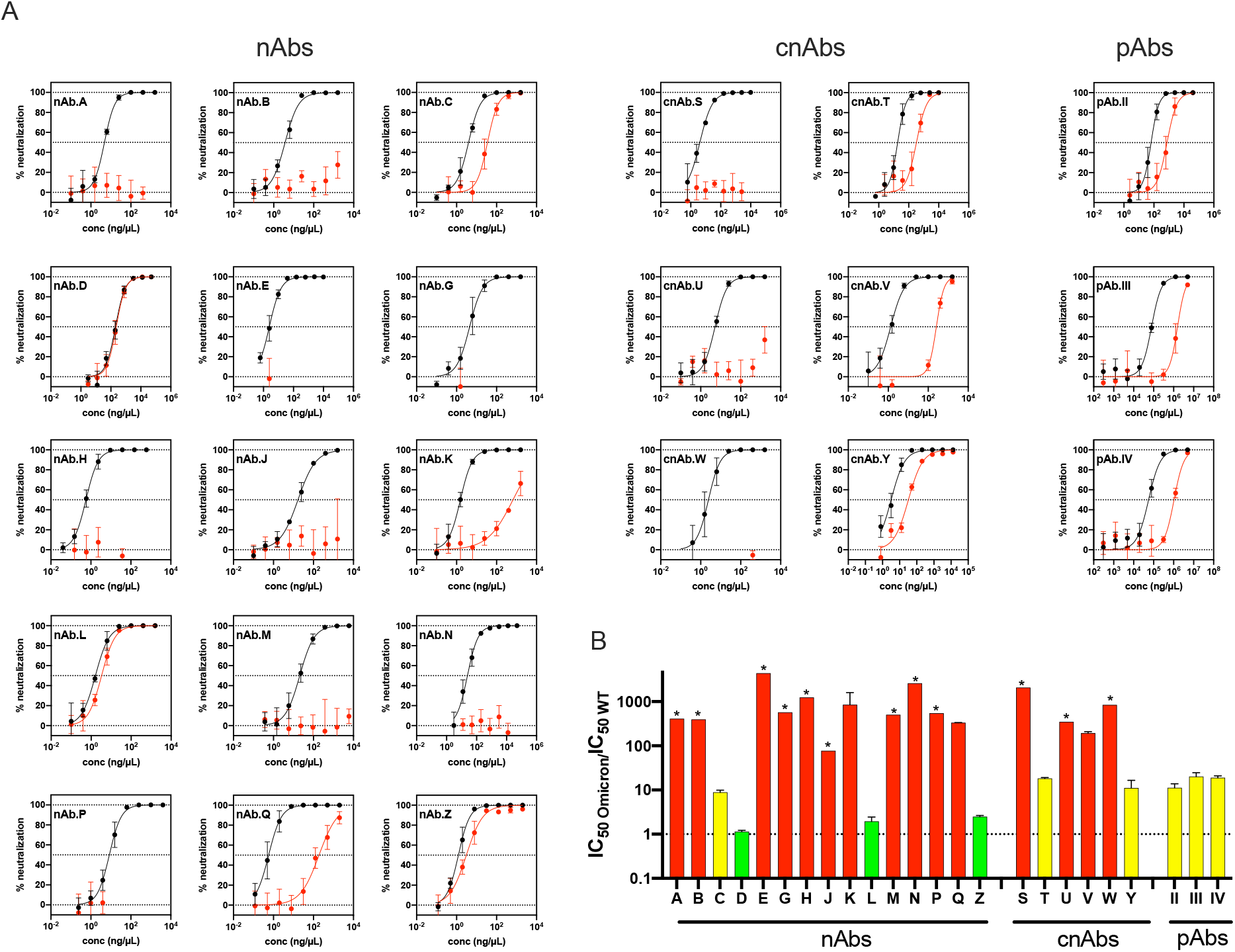
Neutralization of Omicron by therapeutic antibodies. (A) Neutralization curves for each one of the 24 therapeutic antibody products against WT (black) and Omicron (red). (B) Bar graph showing the ratio between the IC_50_ of Omicron and WT for all the antibody products. The sensitivity of the Omicron variant against 15 monoclonal antibodies (nAbs), 6 combination nAbs products (cnAbs), and 3 polyclonal antibodies (pAbs). Red indicates IC_50_ resistance ratios >50, yellow indicates moderate resistance with IC_50_ ratios between 5-50, and green indicates sensitivity comparable to WT with IC_50_ ratios <5. Antibodies for which complete neutralization was not achieved at the highest concentration tested are denoted by *. Data shown represent two independent experiments each with an intra-assay duplicate.

## DISCUSSION

Neutralizing antibodies are widely accepted as an important component of protection against SARS-CoV-2 infection and disease (COVID-19), but efforts to assess antibody titers that correlate with protection are complicated by many factors. These include potential redundancy and synergism of different components of the humoral, cellular, and innate immune system, and differences in variant fitness and host genetics, age, and prior immunity. The risk of infection can also be confounded by human behavior and local public health measures, while measurements of antibody titers can vary with different laboratory methods. Therefore, differences in study populations and laboratory methods are important considerations for assessing the impact of immune evasion by Omicron on medical countermeasures.

Here, using the same lentiviral pseudovirus neutralization platform we measured the change in potency of 24 clinical-stage therapeutic antibodies against Omicron compared to WT (D614G) and compared neutralizing antibodies in sera from two well-characterized cohorts of subjects in prospective clinical studies. Our findings show that most vaccinated individuals have low or undetectable titers against Omicron after the second Pfizer/BNT162b2 vaccination, similar to findings reported by others(*3-7*). However, the third vaccination significantly increased neutralizing titers to levels significantly higher than those elicited by the second vaccination, in agreement with other preliminary studies(*8-14*). It is notable that the post-2^nd^ vaccination and post-3^rd^ vaccination sampling times were similar, indicating that this boost does not simply reflect time since last vaccine. We found no association between sex or age with these neutralizing immune responses, although it is important to note that the study samples were from generally healthy adults, and that the post-vaccine-dose sampling time was overall short (43 ± 17 days). It has been shown that infection followed by vaccination results in neutralizing antibody titers comparable or higher to titers after two vaccinations(*3, 10, 14, 21*). The PASS study included anti-N antibody testing on all blood samples for detection of silent infections. The neutralizing antibody titers among the 17 asymptomatic individuals who seroconverted for N antibodies between the 2^nd^ and 3^rd^ immunization were not higher than those who did not seroconvert. Other studies have shown high antibody titers in convalescent individuals after the 1^st^ or 2^nd^ vaccination(*10, 11*). We did not see an increase after the 3^rd^ vaccination in individuals who seroconverted for N-antibodies. This may be due to reduced antigen load from incident asymptomatic infection or having reached a maximum response after vaccination.

The antigenic cartography analysis suggests that the Omicron variant appears to be antigenically distant from WT and Delta, but this distance seems to decrease after the 3^rd^ vaccination. The apparent broadening of responses to Omicron may be due to boosting of titers to cross-reactive epitopes or improved antibody affinity to shared epitopes, or both. For the convalescent sera, high titers generally correlated with the highest cross-neutralization to Omicron. Continued studies of the breadth against multiple variants and duration of responses after booster vaccinations are urgently needed. In both vaccination and convalescent individuals, our studies suggest that booster vaccinations, even with the original ancestral vaccine antigen, could be beneficial in protecting against Omicron, in agreement with the rapidly accumulating data from many sources(*3, 5, 6*).

Lastly, nine of the fifteen clinical-grade nAbs under EUA or in late stages of clinical development had no measurable IC_50_ against Omicron compared to WT. Also, while most nAbs and cnAbs lost measurable potency, all polyclonal antibodies retained measurable, though reduced with IC_50_. Careful selection of therapeutic antibodies is needed according to variant prevalence. As Omicron continues to acquire additional mutations the products that remain effective could be jeopardized, underscoring the risk associated with this variant and its derivatives.

In summary, our findings indicate that booster doses of mRNA COVID-19 vaccines may afford an increase in protection against Omicron by inducing higher levels of neutralizing antibodies compared to two vaccine doses or the levels of neutralizing antibodies induced by SARS-CoV-2 infection from different variants. The strengths of this study include representation of a broad diversity of genotypes, including within-Delta diversity, comparable sampling times between convalescent and vaccinated subjects, the use of cartography methods, and the availability of subject demographics to interpret how host characteristics may influence Omicron humoral immunity. The limitations of our study include the relatively small numbers of study subjects, restricted timing of sera collection, and the use of a pseudovirus platform as a surrogate to authentic SARS-CoV-2 viruses. Ultimately neutralization titers need to be tied to clinical outcomes. The rapid and unpredictable evolution of SARS-CoV-2 requires continued development and assessments of medical countermeasures.

## MATERIALS AND METHODS

### Vaccinated participants

Details of the PASS study protocol, including the inclusion/exclusion criteria, have been published(*17*). Full details are in the Supplemental Material. Briefly, generally healthy healthcare workers without a history of SARS-CoV-2 infection at screening were enrolled. The study began in August 2020 and involved monthly research clinic visits to obtain serum for longitudinal SARS-CoV-2 antibody testing. The subset of participants included in this study received three doses of Pfizer/BNT162b2 vaccine; none had a PCR-confirmed SARS-CoV-2 infection during follow-up. Participants’ serially-collected serum samples were screened for immunoglobulin G (IgG) reactivity with SARS-CoV-2 spike protein and nucleocapsid protein (N) using a multiplex microsphere-based immunoassay, as described(*22*).

### Unvaccinated infections - study population and general study design

The EPICC study is a cohort study of U.S. Military Health System (MHS) beneficiaries that includes those with a history of SARS-CoV-2 infection, as described previously(*19*). Full details are in Supplemental Material. Enrollment occurred at six Military Treatment Facilities (MTFs). Demographic, comorbidity, COVID illness characteristics, and vaccination status were obtained from the clinical case report form and/or the MHS Data Repository. Biospecimen collection included serial serum samples and upper respiratory specimen swabs.

### Diagnosis of SARS-CoV-2 infection and genotyping of infections used for convalescent sera

SARS-CoV-2 infection was determined by positive clinical laboratory PCR test performed at the enrolling clinical site, or a follow-up upper respiratory swab collected as part of the EPICC study. The specific clinical PCR assay employed at the MTF varied. The follow-up PCR assay (for EPICC specimens) was the SARS-CoV-2 (2019-nCoV) CDC qPCR Probe Assay research use only kits (IDT, Coralville, IA)(*23*); details are in the Supplemental Material. Whole viral genome sequencing was performed on extracted SARS-CoV-2 RNA from PCR-positive specimens; details are in the Supplemental Material. The Pango classification tool (https://cov-lineages.org/) was used for genotype classification (version 3.1.17).

### Ethics

The PASS (IDCRP-126) and EPICC (IDCRP-085) studies were approved by the Uniformed Services University of the Health Sciences Institutional Review Board (IRB) in compliance with all applicable Federal regulations governing the protection of human participants. All PASS and EPICC study participants provided informed consent. The convalescent Beta sera, obtained from a traveler who had moderate-severe Covid-19 in the Republic of South Africa during the peak of the Beta (B.1.351) wave in January 2021, was obtained with informed consent and covered under the US Food and Drug Administration IRB approved expedited protocol # 2021-CBER-045.

### SARS-Cov-2 pseudovirus production and neutralization assays

Lentiviral pseudoviruses were generated and used in neutralization assays, as previously described(*24*). The Omicron spike expression plasmid was generously provided by Nicole Doria-Rose (Vaccine Research Center, National Institutes of Health). Neutralization assays were performed on 293T-ACE2/TMPRSS2 cells stably expressing ACE2 and transmembrane serine protease 2 (BEI # NR-55293). Twenty-four clinical-stage therapeutic antibody products were provided by pharmaceutical companies to support the US government COVID-19 response efforts. The antibody identities are blinded for publications per an agreement with the companies. Neutralization curves were normalized to virus only controls and fitted using nonlinear regression curve (GraphPad Prism, La Jolla, CA). The antibody concentration or sera dilution corresponding to 50% neutralization was defined as NT_50_ for sera or IC_50_ for antibody products. Data reported are averages from at least two independent experiments, each with intra-assay duplicates.

### Antigenic cartography

ACMACS antigenic cartography software (https://acmacs-web.antigenic-441cartography.org/) was used to create a geometric interpretation of neutralization titers against the WT, Delta, and Omicron variants. Each square in the map indicates one antigenic unit, corresponding to two-fold dilution of the antibody in the neutralization assay. Antigenic distance is measured in any direction of the map.

## Supplementary Materials

Supplementary materials and methods

Table S1: SARS-CoV-2 genotypes for the infecting variants

## Acknowledgments

The authors wish to also acknowledge all who have contributed to the EPICC (IDCRP-085) COVID-19 study.

*Brooke Army Medical Center, Fort Sam Houston, TX*: Col J. Cowden; LTC M. Darling; S. DeLeon; Maj D. Lindholm; LTC A. Markelz; K. Mende; S. Merritt; T. Merritt; LTC N. Turner; CPT T. Wellington

*Carl R. Darnall Army Medical Center, Fort Hood, TX*: LTC S. Bazan; P.K Love

*Fort Belvoir Community Hospital, Fort Belvoir, VA:* N. Dimascio-Johnson; MAJ E. Ewers; LCDR K. Gallagher; LCDR D. Larson; A. Rutt

*Henry M. Jackson Foundation, Inc*., *Bethesda, MD:* P. Blair; J. Chenoweth; D. Clark

*Madigan Army Medical Center, Joint Base Lewis McChord, WA*: S. Chambers; LTC C. Colombo; R. Colombo; CAPT C. Conlon; CAPT K. Everson; COL P. Faestel; COL T. Ferguson; MAJ L. Gordon; LTC S. Grogan; CAPT S. Lis; COL C. Mount; LTC D. Musfeldt; CPT D. Odineal; LTC M. Perreault; W. Robb-McGrath; MAJ R. Sainato; C. Schofield; COL C. Skinner; M. Stein; MAJ M. Switzer; MAJ M. Timlin; MAJ S. Wood

*Naval Medical Center Portsmouth, Portsmouth, VA*: S. Banks; R. Carpenter; L. Kim; CAPT K. Kronmann; T. Lalani; LCDR T. Lee; LCDR A. Smith; R. Smith; R. Tant; T. Warkentien

*Naval Medical Center San Diego, San Diego, CA:* CDR C. Berjohn; S. Cammarata; N. Kirkland; D. Libraty; CAPT (Ret) R. Maves; CAPT (Ret) G. Utz

*Tripler Army Medical Center, Honolulu, HI*: S. Chi; LTC R. Flanagan; MAJ M. Jones; C. Lucas; LTC (Ret) C. Madar; K. Miyasato; C. Uyehara

*Uniformed Services University of the Health Sciences, Bethesda, MD*: B. Agan; L. Andronescu; A. Austin; C. Broder; CAPT T. Burgess; C. Byrne; COL K Chung; J. Davies; C. English; N. Epsi; C. Fox; M. Fritschlanski; M. Grother; A. Hadley; COL P. Hickey; E. Laing; LTC C. Lanteri; LTC J. Livezey; A. Malloy; R. Mohammed; C. Morales; P. Nwachukwu; C. Olsen; E. Parmelee; S. Pollett; S. Richard; J. Rozman; J. Rusiecki; E. Samuels; M. Sanchez; A. Scher; CDR M. Simons; A. Snow; K. Telu; D. Tribble; L. Ulomi

*United States Air Force School of Medicine, Dayton, OH*: SSgt T. Chao; R. Chapleau; M. Christian; A. Fries; C. Harrington; S. Huntsberger; K. Lanter; E. Macias; J. Meyer; S. Purves; K. Reynolds; J. Rodriguez; C. Starr

Womack Army Medical Center, Fort Bragg, NC: B. Barton; LTC D. Hostler; LTC J. Hostler; MAJ K. Lago; C. Maldonado; J. Mehrer

*United States Army Medical Research Institute of Infectious Diseases, Frederick, MD*: MAJ J. Kugelman

*William Beaumont Army Medical Center, El Paso, TX*: MAJ T. Hunter; J. Mejia; R. Mody; R. Resendez; P. Sandoval; M. Wayman

*Walter Reed National Military Medical Center, Bethesda, MD:* I. Barahona; A. Baya; A. Ganesan; MAJ N. Huprikar; B. Johnson

*Walter Reed Army Institute of Research, Silver Spring, MD*: S. Peel

We also thank Douglas Pratt and Hector Izurieta (US Food and Drug Administration) for critical review of the manuscript.

## Author contributions

Conceptualization: SL, SDP, RV, EG, CCB, LB, KJE, EDL, THB, EM, CDW Methodology: SL, SDP, SNN, WW, RV, ACF, EG, SAC, CCB, LB, KJE, EDL, LCK, EM, CDW

Investigation: SL, SDP, SNN, WW, RW, RV, NJE, ACF, DAL, CJC, RM, AG, EG, SAC, GW, LB, KJE, ECE, TL, CDW

Visualization: SL, SDP, WW, RV, LCK, CDW

Funding acquisition: SDP, AEE, DRT, THB, EM, CDW

Project administration: SDP, RV, BKA, MH-P, MPS, DRT, AEE, KJE, EDL, THB, EM, CDW Supervision: SDP, CCB, EDL, EM, THB, CDW

Writing – original draft: SL, SDP, CDW

Writing – review & editing: SL, SDP, SNN, WW, RW, RV, NJE, ACF, BKA, DAL, CJC, RM, ECE, TL, AG, EG, MH-P, SAC, MPS, LCK, GW, CCB, DRT, LB, AEE, KJE, EDL, THB, EM, CDW

## Disclaimer

E. Laing, T. Burgess, D. Tribble, A Fries, C. Broder, E. Mitre, M. Hollis-Perry, S. Lusvarghi, S. Pollett, SN. Neerukonda, W. Wang, R. Wang, R. Vassell, LC. Katzelnick, L Bentley, AE. Eakin, KJ. Erlandson and CD. Weiss are service members or employees of the U.S. Government. This work was prepared as part of their official duties. Title 17 U.S.C. §105 provides that ‘Copyright protection under this title is not available for any work of the United States Government.’ Title 17 U.S.C. §101 defines a U.S. Government work as a work prepared by a military service member or employee of the U.S. Government as part of that person’s official duties.

The contents of this publication are the sole responsibility of the author (s) and do not necessarily reflect the views, opinions, or policies of Uniformed Services University of the Health Sciences (USUHS); the Department of Defense (DoD); the Departments of the Army, Navy, or Air Force; the Defense Health Agency, Brooke Army Medical Center; Walter Reed National Military Medical Center; Naval Medical Center San Diego; Madigan Army Medical Center; United States Air Force School of Aerospace Medicine; Fort Belvoir Community Hospital; the Henry M. Jackson Foundation for the Advancement of Military Medicine Inc; the National Institutes of Health; the Office of the Assistance Secretary for Preparedness and Response, US Department of Health and Human Services (HHS); the Biomedical Advanced Research and Development Authority, HHS; or the US Food and Drug Administration. Mention of trade names, commercial products, or organizations does not imply endorsement by the U.S. Government. The investigators have adhered to the policies for protection of human subjects as prescribed in 45 CFR 46.

## Group authors EPICC and PASS (MEDLINE indexed)

Brooke Army Medical Center, Fort Sam Houston, TX: Katrin Mende

Clinical Trials Center, Infectious Diseases Directorate, Naval Medical Research Center, Silver Spring, MD, USA: Christopher Duplessis, Kathleen F. Ramsey, Anatalio E. Reyes, Yolanda Alcorta, Mimi A. Wong, Santina Maiolatesi, Keishla Morales Padilla

Fort Belvoir Community Hospital, Fort Belvoir, VA: Derek Larson

Madigan Army Medical Center, Joint Base Lewis McChord, WA: Rhonda E. Colombo

Naval Medical Center Portsmouth, Portsmouth, VA: Alfred Smith

Uniformed Services University of the Health Sciences, Bethesda, MD: Stephanie Richard, Julia

Rozman, Alyssa R. Lindrose, Matthew Moser, Belinda Jackson-Thompson, Margaret Sanchez-Edwards, Luca Illinik, Julian Davies, Orlando Ortega, Edward Parmelee

United States Army Medical Research Institute of Infectious Diseases, Frederick, MD: MAJ J. Kugelman

Walter Reed National Military Medical Center, Bethesda, MD: Nikhil Huprikar

