## supplemental table for "SARS-CoV-2 Omicron neutralization by therapeutic antibodies, convalescent sera, and post-mRNA vaccine booster"

**Supplementary Table. SARS-CoV-2 genotypes for the infecting variants.**

| Sample # | Pangolin 3.1.17 (2021-12-06) | Spike Substitutions | Spike Deletions |
|---|---|---|---|
| Conv-01 | AY.74 | S:T19R,S:G142D,S:R158G,S:A222V,S:L452R,S:T478K,S:D614G,S:P681R,S:D950N | S:E156-,S:F157- |
| Conv-02 | AY.25 | S:T19R,S:S112L,S:G142D,S:R158G,S:L452R,S:T478K,S:D614G,S:P681R,S:D950N | S:E156-,S:F157- |
| Conv-03 | AY.47 | S:T19R,S:G142D,S:R158G,S:A222V,S:V289I,S:L452R,S:T478K,S:D614G,S:P681R,S:D950N | S:E156-,S:F157- |
| Conv-04 | B.1.617.2 | S:T19R,S:K77T,S:G142D,S:R158G,S:G181V,S:L452R,S:T478K,S:D614G,S:A653V,S:P681R,S:D950N | S:E156-,S:F157- |
| Conv-05 | B.1.617.2 | S:T19R,S:K77T,S:G142D,S:R158G,S:G181V,S:L452R,S:T478K,S:D614G,S:A653V,S:P681R,S:D950N | S:E156-,S:F157- |
| Conv-06 | AY.14 | S:T19R,S:G142D,S:R158G,S:L452R,S:T478K,S:D614G,S:P681R,S:D950N | S:E156-,S:F157- |
| Conv-07 | AY.14 | S:T19R,S:G142D,S:R158G,S:L452R,S:T478K,S:D614G,S:P681R,S:D950N | S:E156-,S:F157- |
| Conv-08 | AY.25 | S:T19R,S:S112L,S:G142D,S:R158G,S:L452R,S:T478K,S:D614G,S:P681R,S:D950N | S:E156-,S:F157- |
| Conv-09 | B.1.617.2 | S:T19R,S:T95I,S:G142D,S:R158G,S:L452R,S:T478K,S:D614G,S:P681R,S:D950N,S:G1124V | S:E156-,S:F157- |
| Conv-10 | AY.62 | S:T19R,S:G142D,S:R158G,S:A222V,S:L452R,S:T478K,S:D614G,S:P681R,S:G946V,S:D950N | S:E156-,S:F157- |
| Conv-11 | AY.25 | S:T19R,S:G142D,S:R158G,S:L452R,S:T478K,S:D614G,S:P681R,S:D950N | S:E156-,S:F157- |
| Conv-12 | AY.44 | S:T19R,S:T22I,S:G142D,S:R158G,S:L452R,S:T478K,S:D614G,S:P681R,S:D950N | S:E156-,S:F157- |
| Conv-13 | AY.119 | S:T19R,S:T95I,S:G142D,S:R158G,S:L452R,S:T478K,S:D614G,S:P681R,S:D950N | S:E156-,S:F157- |
| Conv-14 | B.1.617.2 | S:T19R,S:G142D,S:R158G,S:L452R,S:T478K,S:D614G,S:P681R,S:D950N | S:E156-,S:F157- |
| Conv-15 | B.1.617.2 | S:T19R,S:G142D,S:R158G,S:L452R,S:T478K,S:D614G,S:P681R,S:D950N,S:V1264L | S:E156-,S:F157- |
| Conv-16 | B.1.617.2 | S:T19R,S:T95I,S:G142D,S:R158G,S:L452R,S:T478K,S:D614G,S:P681R,S:D950N | S:E156-,S:F157- |
| Conv-17 | B.1.617.2 | S:T19R,S:G142D,S:M153I,S:R158G,S:A222V,S:L452R,S:T478K,S:D614G,S:P681R,S:D950N | S:E156-,S:F157- |
| Conv-18* | B.1.351 | | |
| Conv-19 | B.1.351 | S:D80A,S:D215G,S:K417N,S:E484K,S:N501Y,S:D614G,S:A701V | |
| Conv-20 | B.1 | S:D614G | |
| Conv-21 | B.1 | S:D614G | |
| Conv-22 | B.1 | S:D614G | |
| Conv-23 | B.1 | S:D614G | |
| Conv-24 | B.1 | S:D614G | |
| Conv-25 | B.1 | S:D614G | |
| Conv-26 | B.1 | S:D614G | |
| Conv-27 | B.1 | S:D614G | |
| Conv-28 | B.1 | S:D614G | |
| Conv-29 | B.1 | S:D614G | |
| Conv-30 | B.1.1.7 | S:N501Y,S:A570D,S:D614G,S:P681H,S:T716I,S:S982A,S:D1118H | S:H69-,S:V70-,S:Y144- |
| Conv-31 | B.1.1.7 | S:N501Y,S:A570D,S:D614G,S:P681H,S:T716I,S:S982A,S:D1118H | S:H69-,S:V70-,S:Y144- |
| Conv-32 | B.1.1.7 | S:N501Y,S:A570D,S:D614G,S:P681H,S:T716I,S:S982A,S:D1118H,S:K1191N | S:H69-,S:V70-,S:Y144- |
| Conv-33 | B.1.1.7 | S:N501Y,S:A570D,S:D614G,S:P681H,S:T716I,S:S982A,S:D1118H | S:H69-,S:V70-,S:Y144- |
| Conv-34 | B.1.1.7 | S:N501Y,S:A570D,S:D614G,S:P681H,S:T716I,S:S982A,S:D1118H | S:H69-,S:V70-,S:Y144- |
| Conv-35 | B.1.2 | S:D614G | |
| Conv-36 | B.1.2 | S:G257D,S:D614G | |
| Conv-37 | B.1.2 | S:D614G | |
| Conv-38 | B.1.2 | S:D614G | |
| Conv-39 | B.1.2 | S:D614G | |
| Conv-40 | B.1.2 | S:D614G | |

*The sequencing information from this individual was not avalable (see methods)
