## Supplementary methods for "SARS-CoV-2 Omicron neutralization by therapeutic antibodies, convalescent sera, and post-mRNA vaccine booster"

### MATERIALS AND METHODS

#### Vaccinated participants

Details of the Prospective Assessment of SARS-CoV-2 Seroconversion (PASS) study protocol, including details of the inclusion/exclusion criteria, have been published (*17*). Inclusion criteria included being generally healthy, ≥ 18 years old, and employed at the WRNMMC. Exclusion criteria included history of COVID-19, IgG seropositivity for SARS-CoV-2, and being severely immunocompromised at time of screening. The study was initiated in August 2020, with rolling enrollment and monthly research clinic visits to obtain serum for longitudinal SARS-CoV-2 antibody testing. The subset of participants included in this study received two doses of Pfizer/BNT162b2 vaccine by January 26, 2021 and had no serological or PCR evidence of SARS-CoV-2 infection prior to two doses of vaccine. These subjects received a 3$^{rd}$ dose of Pfizer/BNTech162b2 vaccine by Nov 18, 2021. No subject included in this analysis had a clinically apparent PCR-confirmed SARS-CoV-2 infection during follow-up. Participants' serum samples were collected monthly through September of 2021, and then quarterly, diluted 1:400 and 1:8000, and screened for immunoglobulin G (IgG) reactivity with SARS-CoV-2 spike protein and nucleocapsid protein (N), and four human coronavirus (HCoV) spike proteins using a multiplex microsphere-based immunoassay, as previously described. For this study sample, we selected individuals with sera available at 3 to 6 weeks after symptom onset, with complete or near complete and high coverage spike gene sequence availability, with no prior vaccination, and with a diverse set of genotypes including Delta and non-Delta variants.

**Unvaccinated infections - study population and general study design**

The Epidemiology, Immunology, and Clinical Characteristics of Emerging Infectious Diseases with Pandemic Potential (EPICC) study is a cohort study of U.S. Military Health System (MHS) beneficiaries which includes those with a history of SARS-CoV-2 infection(*19*). Eligibility criteria for enrollment included those presenting to clinical care with COVID-19-like illness and for SARS-CoV-2 polymerase chain reaction (PCR) testing. The EPICC study has been enrolling since March 2020. For this sample set, EPICC enrollment occurred at six Military Treatment Facilities (MTFs): Brooke Army Medical Center, Fort Belvoir Community Hospital, Madigan Army Medical Center, Naval Medical Center Portsmouth, Walter Reed National Military Medical Center, and the William Beaumont Army Medical Center.

Study procedures for these subjects with SARS-CoV-2 infection included collection of demographic data and completion of a clinical case report form (CRF) to characterize the acute COVID-19 illness. Biospecimen collection included serial serum samples for immune response analysis, and upper respiratory specimen swabs for virological analysis. For all enrolled subjects, we also abstracted MHS-wide healthcare encounter data from the Military Health System Data Repository (MDR) to determine comorbidities. Vaccination status was ascertained by the MDR record in addition to the CRF and questionnaire self-report.

**Diagnosis of SARS-CoV-2 infection and genotyping of infections used for convalescent sera**

SARS-CoV-2 infection was determined by positive PCR clinical laboratory test performed at the enrolling clinical site, or a follow-up upper respiratory swab collected as part of the EPICC study. The specific PCR assay employed at the MTF varied. The follow-up PCR assay (used for

EPICC specimens) was the SARS-CoV-2 (2019-nCoV) CDC qPCR Probe Assay research use only kits (IDT, Coralville, IA)(*22*). This assay targets two regions of the SARS-CoV-2 nucleocapsid (N) gene (N1 and N2), with an additional control target to detect the human RNase P gene (RP). We considered a positive SARS-CoV-2 infection as positive based on a cycle threshold value of less than 40 for both N1/N2 gene targets.

Whole viral genome sequencing was performed on extracted SARS-CoV-2 RNA from PCR positive specimens. A 1200bp amplicon tiling strategy was used (dx.doi.org/10.17504/protocols.io.btsrnnd6). Amplified product was prepared for sequencing using NexteraXT library kits (Illumina Inc., San Diego, CA). Libraries were run on the Illumina NextSeq 550 sequencing platform. BBMap v. 38.86 and iVar v. 1.2.2 tools were used for genome assembly. The Pango classification tool (https://cov-lineages.org/) was used for genotype classification (version 3.1.17).

The infecting genotype for one subject (Conv-19, Supplementary Table) was determined from a viral sequence derived from an alternative sequencing platform (Illumina MiSeq). Briefly, cDNA synthesis was performed with the Superscript IV first-strand synthesis system (Life Technologies/Invitrogen, Carlsbad, CA). Multiplex PCR was performed with the ARTIC v3 primer set, designed to amplify overlapping regions of the Sars-CoV-2 reference genome (MN908947.3). Primer and genomic alignment position information is available here: https://github.com/artic-network/artic-ncov2019/tree/master/primer_schemes/nCoV-2019/V1. PCR products were purified with the MinElute PCR purification kit (QIAgen, Valencia, CA). Libraries were prepared with the SMARTer PrepX DNA Library Kit (Takara Bio, Mountain View, CA), using the Apollo library prep system (Takara Bio, Mountain View, CA). The libraries were evaluated for quality using the Agilent 2200 TapeStation (Agilent, Santa Clara, CA). After quantification by real-time PCR with the KAPA SYBR FAST qPCR Kit (Roche, Pleasanton, CA), libraries were diluted to 10 nM.
